# Causal inference using deep-learning variable selection identifies and incorporates direct and indirect causalities in complex biological systems

**DOI:** 10.1101/2021.07.17.452800

**Authors:** Zhenjiang Fan, Kate F. Kernan, Panayiotis V. Benos, Scott W. Canna, Joseph A. Carcillo, Soyeon Kim, Hyun Jung Park

## Abstract

In complex diseases, causal structure learning across biological variables is critical to identify modifiable triggers or potential therapeutic agents. A limitation of existing causal learning methods is that they cannot identify indirect causal relations, those that would interact through latent mediating variables. We developed the first computational method that identifies both direct and indirect causalities, causal inference using deep-learning variable-selection (causalDeepVASE). To accurately identify indirect causalities and incorporate them with direct causalities, causalDeepVASE develops a deep neural network approach and extends a flexible causal inference method. In simulated and biological data of various contexts, causalDeepVASE outperforms existing methods in identifying expected or validated causal relations. Further, causalDeepVASE facilitates a systematic understanding of complex diseases. For example, causalDeepVASE uniquely identified a possible causal relation between IFNγ and creatinine suggested in a polymicrobial sepsis model. In future biomedical studies, causalDeepVASE can facilitate the identification of driver genes and therapeutic agents.

## INTRODUCTION

In complex biological systems such as humans, molecular and clinical features demonstrate variable associations in diseases such as cancer, asthma, and sepsis (1)(2)(3). Identifying the causal relations in the association is critical to improving translational understanding since causal relations can be used to identify potential therapeutic agents that can control the diseases. For example, if an abnormal expression of certain genes plays causative roles in developing a disease, then further efforts could be made to use these genes for diagnostics and control these genes to treat the disease. Bioinformatic methods have been developed to identify the causal structure from associations between variables. Among the methods, causalMGM (causal mixed graphical model) and the Degenerated Gaussian (DG) score have been shown to achieve great sensitivity and specificity in simulated and biological data and can handle mixed types of variables (4)(5). CausalMGM first identifies variable associations by testing the significance of the interaction terms in a pseudo-likelihood function (mixed graphical model (MGM)) (6). From the variable associations, it learns the causal structure using the constraint-based PC (Peter and Clark) algorithm or its variants (e.g., PC-Stable (7), FCI-MAX (8), etc.). Especially, MGM-FCI-MAX can learn a causal graph in a dataset with latent confounders. The other method, DG, does not directly identify variable associations, but, given variable associations, learns the causal structure by evaluating the direction of the interaction terms in a conditional Gaussian likelihood score with certain assumptions (see Methods).

However, both methods share a critical limitation in learning causalities in complex biological systems. Although they can identify direct associations between variables by testing their interaction, they cannot identify indirect associations that would interact through latent mediating variables. Recent studies show that complex biological systems with multiple biological layers (such as transcriptome, methylome, and proteome) can have indirect relationships between variables through mediators in the multiple layers (9), illustrating the necessity to consider not only direct but also indirect associations in causal structure learning. For example, in childhood asthma, 366 (88%) of 418 eQTL SNPs affect gene expression levels both directly and indirectly through DNA methylation (10). Therefore, to learn comprehensive causalities in complex biological systems such as asthma, it is critical to identify both direct and indirect causalities.

In this manuscript, we developed one of the first computational methods that identifies direct and indirect causalities among the input variables, causal inference using deep-learning variable selection (causalDeepVASE). This novel method decomposes the task of learning causalities into three steps: identifying direct and indirect associations and learning causalities from both types of associations. In the first step, to identify direct associations, we employed the MGM approach implemented in causalMGM due to its validated performance (11). In the second step, to identify indirect associations, we utilized a deep-learning approach by modelling the mediating layers with perceptron layers (layers of linear classifiers) and inferring their interactions through computational optimization. In addition, we use causalDeepVASE to estimate the effect size of the associations using knockoff variables (12) as the negative control. Knockoff variables are to mimic the correlation structure of the input variables in a very specific way that will allow for effect size estimation and FDR control of the corresponding input variables. Estimating the effect size can facilitate a translatable understanding of the causal structure since downstream experiments or clinical trials can target only a limited number of strong causalities for technical and practical limitations. Using knockoff variables as the negative control allowed for successful estimation of the effect size of important variables in Deep feature selection using Paired-Input Nonlinear Knockoffs (deepPINK) (13). This approach will successfully estimate the effect size of indirect associations.

In the third step, we address the main challenge to learn causalities from both direct and indirect associations, as such associations were not previously considered. To develop a causality learning approach for this purpose, we considered two conditions, decomposability and flexibility. First, the optimal causal structure based on any novel approach must be decomposable to the optimal causal relation in each association. This decomposability will allow us to determine the causal relation of each association without referring to other associations, whether it is a direct or indirect association. Second, the approach needs to be flexible in learning causalities generated outside of the assumed class. This flexibility will allow us to hypothesize that it can learn causalities from indirect associations. Identifying that the Degenerate Gaussian (DG) score satisfies these conditions with certain assumptions, we extended DG to learn causalities in causalDeepVASE. Altogether, by identifying and incorporating indirect associations and extending DG to learn indirect causality, causalDeepVASE increases the power to learn causal relations in complex biological systems compared to existing methods that only identify direct causalities.

Previously, deep-learning approaches have been applied to learn causal relations in biological and clinical data. However, causalDeepVASE is the first one utilizing a deep-learning approach to learn indirect causal relations. For example, Yu et al. used a deep-learning generative model to search for optimal causal relations under an optimization criterion (14). Also, Young et al. used a deep-learning approach to model latent mediators between cause and the effect variables (15). Although Young et al.’s approach is close to our design, the difference is that they assume to know which of the input variables are the cause and which are the effect variables. On the other hand, causalDeepVASE does not assume to know the cause and the effect variables and identify the cause-and-effect relationship in the data.

## MATERIAL AND METHODS

### Biological data

TCGA breast invasive carcinoma (BRCA) data in this work were downloaded from https://tcga.xenahubs.net. It consists of the gene expression RNAseq dataset (dataset ID: TCGA.BRCA.sampleMap/HiSeqV2) and the clinical phenotype dataset (dataset ID: TCGA.BRCA.sampleMap/BRCA_clinicalMatrix). For the gene expression dataset, we selected 500 expressed genes based on their variances and added another gene ERBB2 to the selected gene set. For the clinical dataset, we used 10 well-known clinical status features: PAM50Call_RNAseq, HER2_Final_Status_nature2012, Converted_Stage_nature2012, Node_nature2012, breast_carcinoma_progesterone_receptor_status, breast_carcinoma_estrogen_receptor_status, lymph_node_examined_count, person_neoplasm_cancer_status, pathologic_stage; and then we only kept 601 samples who have no missing values in the 10 selected features.The body-mass index (BMI) data has 90 samples and 301 features, where 214 of these 301 features micronutrients and the rest 87 of them are bacteria genera. We used the same data pre-processing procedure as (13) used in their work. The pediatric sepsis data contains 283 septic patients and 56 clinical and laboratory features. The data were collected from 9 PICUs in the Eunice Kennedy Shriver National Institutes of Child Health and Human Development Collaborative Pediatric Critical Care Research Network (including Children’s Hospital of Pittsburgh, Children’s Hospital of Philadelphia, Children’s National Medical Center, Children’s Hospital of Michigan, Nationwide Children’s Hospital, Children’s Hospital of Los Angeles, St. Louis Children’s Hospital, C. S. Mott Children’s Hospital, and Mattel Children’s Hospital at the University of California Los Angeles) (16). Another work analyzing this data is in progress. This data is currently not deposited in the public domain yet.

### causalDeepVASE

To select variables linearly and non-linearly associated with each of the input variables, causalDeepVASE develops regularized linear and non-linear model. For linear model, causalDeepVASE identifies associated variables in a pairwise Markov Random Field or undirected graphical model. Recently, Lee and Hastie proposed the log-likelihood of graphical model Θ of M continuous variables (following normal distribution, denoted by *x* ∈ *X*) and N categorical variables (denoted by *y* ∈ *Y*) as follows.

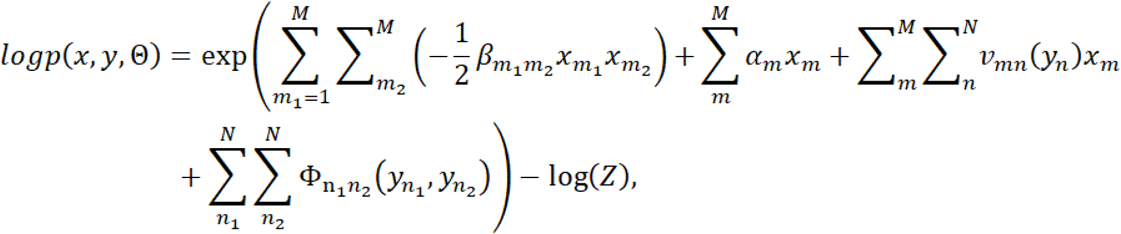

where *β*_*m*_1_ *m*_2__ is the interaction between two continuous variables, *x*_*m*_1__ and *x*_*m*_2__, *α*_*m*_ is the potential of continuous variable *x*_m_, *v_mn_* is a matrix of interaction parameters between the continuous variable *x_m_* with each index of the categorical variable *y_n_*, Φ_n_1_n_2__ is a matrix of interaction parameters between the discrete variable *y*_*n*_1__ and *y*_*n*_2__ (indexed by their levels) (6). While calculating the partition function *Z* can be expensive, it is possible to optimize the log-likelihood edge by edge (5). Overall, this equation models the log-likelihood of interactions of continuous variables and categorical variables as a multinomial linear regression. To ensure sparsity and select associated variables in the regression model, Sedgewick et al. introduced data sparsity penalties for interactions between continuous, categorical, and across continuous and categorical variables (*λ_cc_*, *λ_cd_*, *λ_dd_*, respectively) as follows.

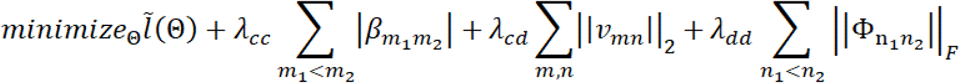

To identify non-linear variable associations, causalDeepVASE sets multiple perceptron layers to model the multiple hidden layers possibly operating on the associated variables. To address the second consideration and select variables *X*′ ⊂ *X_\m_* for *x_m_*, causalDeepVASE considers model-X knockoff variable for each *x_n_* ∈ *X_/m_* according to the following definition (17).

#### Definition 1.

Model-X knockoff variables for the family of random variables 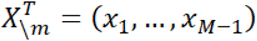 are a new family of random variables 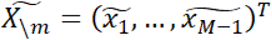 that satisfies two properties: (1) 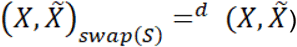 for any subset *S* ⊂ {1,…, *M* – 1}, wheie *swap*(*S*) means swapping *x_j_* and 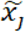 for each *j* ∈ *S* and =^*d*^ denotes equal in distribution, and (2) 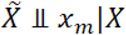, that is, 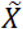 is independent of response *x_m_* given variable *X*. By pairing knockoff variables 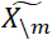 with the corresponding input variables in *X_\m_* in the multiple layers of perceptrons for each response *x_m_* and running optimization concerning *x_m_*, one can quantify strong variables based on 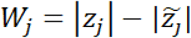, where *z_j_*, and 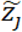 indicate the node weight determined in the optimization. Based on *W_j_*, causalDeepVASE determines the associated variables (*x_m_*, *x_n_* ∈ *X_\m_*) using stopping criteria for the user-specific FDR level *q* as follows ^31^. When causalDeepVASE embeds this process in our module 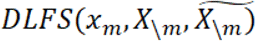, it identifies all non-linearly associated variables as 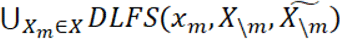.

Also, while the derivation above considers continuous variables, causalDeepVASE transforms categorical variable *x_n_* with *p* categories into a set of *p* random variables that can be taken as continuous as follows. *x_n_* = *I*_1_(*x_n_*),…, *I_p_*(*x_n_*), where *I*_1_(*x_n_*) is the indicator function such that *I_j_*(*x_n_*) = 1 if *x_n_* = *j* and *I_j_*(*x_n_*) = 0 otherwise (one-hot vector). Since this transformation originally leads to a degenerate distribution that supports only space with one less dimension, causalDeepVASE drops the last indicator variable in each one-hot vector as suggested. Then, causalDeepVASE ensures to produce a directed acyclic graph (DAG) by algorithmically removing the causal relations that create the cycle. If there are multiple causal relations, the removal of which will lead to a DAG, causalDeepVASE removes the one with the least effect size.

To estimate the degree of the association between variable *X* and *Y*, the regression model typically tests conditional dependence of them given a conditioning set of variables *S*. If *X* and *Y* are independent given *S*(*X* ⊥ *Y*|*S*), then the following equation would hold.

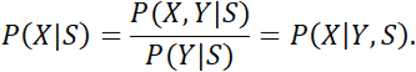

So, to test the independence of *X* and *Y* given *S*, it suffices to test if *P*(*XS*) = *P*(*X*|*Y*, *S*) which can be done via likelihood ratio test (LRT) of two regression models that model *X* given *S* with and without *Y*. Since this is known to follow the chi-squared distribution, causality is learned based on the chisquare test formulated as follows.

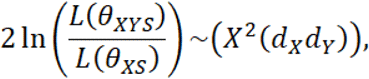

where *θ* represents the regression coefficients of the corresponding model.

### Degenerate Gaussian score

Assume a dataset contains *p* random variables *X* = (*X*_1_, *X*_2_,…, *X*_2_) where the dataset is generated from a causal mechanism. Generally, this causal mechanism is represented using a directed acyclic graph (DAG) *G*. *G* consists of a set of numeric vertices *V* and a set of directed edges *E*, where the vertices *V* are the numeric representations of the random variable set *X* and *E* are a set of edges that link the vertices in *V*. If vertex *V_i_* causes another vertex *V_j_*, *V_i_* a parent of *V_j_* and *V_j_* is a child of *V_j_*, which is denoted as *V_j_* → *V_j_*. *V_j_* may have multiple parents, which is denoted as 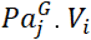 may have multiple children, which is denoted as 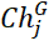. Intuitively, the distribution of variable *X_j_* is impacted by its parents in a DAG or by nothing if it does not have any parents, which can be defined as 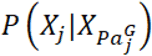. In other words, it should be conditionally independent of other vertices given its parents according to Markov’s condition and d-separation theory. The joint distribution of all the vertices in a DAG would be defined as the product of the distribution of each vertex in it:

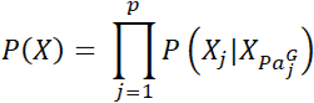

If we log transform the above equation, we get:

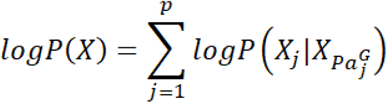

The transformation above enables us to score subgraphs of the original DAG as the original equation naturally decomposes into sub-problems. DG utilizes this decomposability attribute as its score criteria to locally score subgraphs to find an optimal causal structure:

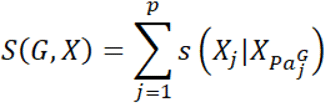

Score-based algorithms can use it because of the following reasons: 1) it is decomposable since those algorithms greedily search and score the subgraph space; 2) it gives the same score while scoring any two Markov equivalent DAGs; 3) it can consistently produce correct scores.

The first step of DG score is to embeds all the discrete variables into a continuous space. For each variable *X_j_*, DG score checks to what data type it belongs, discrete or continuous. Do nothing if it is a continuous variable. If it is a discrete variable, DG score embeds it into a continuous space using their one-hot vector representations as the following equation does:

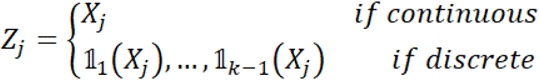

where 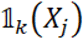 is the indicator function that creates one-hot vector representations for a discrete variable (in each new vector 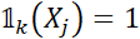 if *X_j_* = *k* and 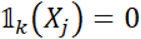 otherwise).

For the newly generated dataset, a full covariance matrix is computed as the covariance matrix will be used for calculating the Gaussian log-likelihood. With this new dataset, the DG score is defined as:

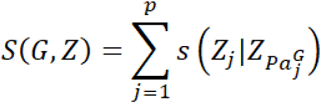

where

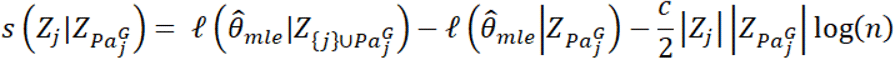

And *c* is a penalty discount used to tune the density of the resulting graph. 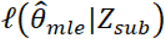, which is the log-likelihood of a subset of Z, is computed using the Gaussian log-likelihood function: where 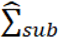 is the partial covariance matrix for the variables in *sub*.

### Single Index model

The nonlinear simulation datasets were generated using a single index model, which is widely used in economics, statistics, and machine learning. Each simulation dataset consists of two parts: a response variable 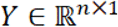; a set of independently and identically distributed random variables 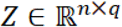 which have no associations with *Y*; and a set of independently and identically distributed random variables 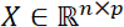 which have a nonlinear association with *Y*. The following model was used for generating these nonlinear datasets:

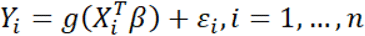

where *g* is an unknown link function, *Y_i_* is the *ith* response value, *X_i_* is the feature vector corresponding to the *ith* observation, and *ε_i_* is the noise added to the *ith* response variable. The matrix *Z* and *X* together was simulated independently from a distribution 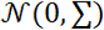 with a precision matrix ∑^-1^ = (*ρ*^|*j*-*k*|^)_1≤*j*, *k*≤(*q*+*p*)_ with *ρ* = 0.5. The distribution for noise *ε* was simulated from 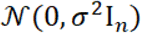, where *σ* is set as 1.

### Pre- and post-processing

To reduce false-positive discoveries, causalDeepVASE carries out several pre- and post-processing steps. As a preprocessing step, causalDeepVASE filters out variable pairs that are conditionally independent on all the other variables since their surficial association must represent their causal relations to the variables the independence is conditioned on, not to each other. Thus, we filtered those whose inverse covariance value is less than 0.0001 after calculating the value using scipy.linalg.inv. Also, as an optional postprocessing step, causalDeepVASE can detect cycle components among the associated variables, so users can refine an acyclic graph with their prior knowledge.

To investigate the dietary effect on the human gut microbiome, we downloaded the nutrient intake/bacteria genera and BMI data from deepPINK site that has 214 micronutrients and 87 genera. For a consistent computation using both the data, the nutrient values are normalized using the residual method to adjust for caloric intake and then standardized (18). Furthermore, the genera data are extracted using 16S rRNA sequencing from stool samples. This data is first log-ratio transformed to get rid of the sum-to-one constraint and then centralized. Following (18), 0s are replaced with 0.5 before converting the data to a compositional form. With both the nutrient intake and genera composition as predictors, we treat BMI as the response.

## RESULTS

### <u>causal</u> inference using <u>deep</u>-learning <u>va</u>riable <u>se</u>lection (causalDeepVASE)

Given biomedical data of mixed types (**Figure 1A**), causalDeepVASE exploits not only direct (**Figure 1B**) but also indirect associations (**Figure 1C**). For this, causalDeepVASE formulated this problem as a set of variable selection problems; it sets each variable as response and selects the other variables related to the response variable. To identify variables directly related to the response variable, causalDeepVASE develops a penalized regression function with the interaction terms connecting the response variable and each of the other variables and maximizes the likelihood with sparsity penalties (**Figure 1B1**). This approach was implemented in causalMGM (11) and showed a good performance at identifying direct associations in simulated and biological data. To identify variables indirectly related to the response variable, it is difficult to use a statistical model, since it requires identifying and incorporating all variables mediating the indirect associations in the model. Instead, causalDeepVASE incorporates a deep learning approach, where multiple perceptron layers are put between the response variable and all the other variables to select the variables that function as mediators (**Figure 1C1**). Another novelty of causalDeepVASE is the uniform estimation of the effect size across direct and indirect associations. To this end, the input variables are paired with the corresponding knockoff variables. Since knockoff variables mimic the arbitrary dependence structure among the input variables while conditionally independent from the response given the input variables (12), this paring allows estimating the strength of each input variable in reference to the corresponding knockoff variables.

**Figure 1.**
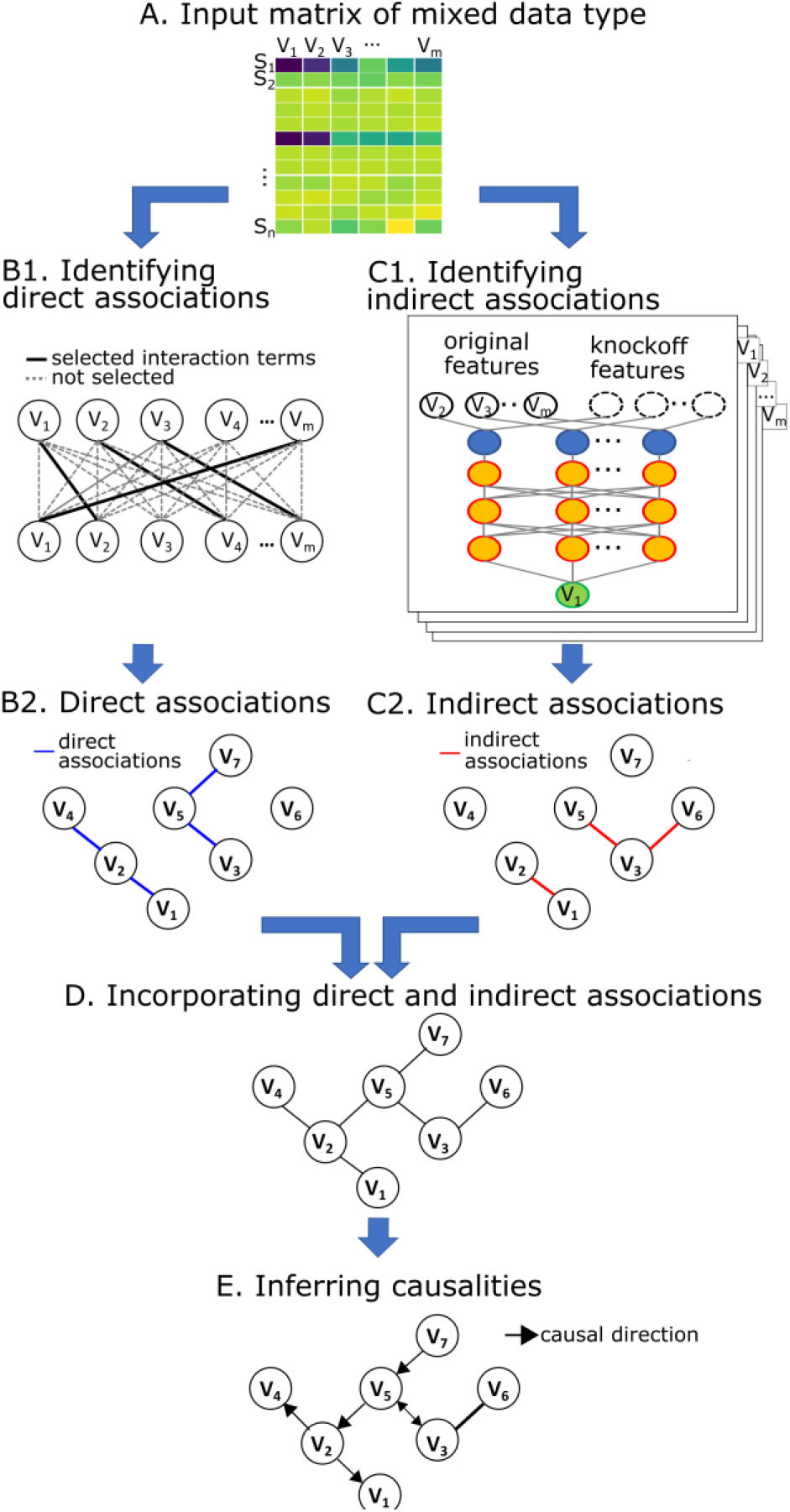
Overview of causalDeepVASE. **(A) An input data matrix** consisting of m variables, either molecular (transcriptomic, epigenetic, genotypic) and clinical variables, either continuous or ordinal categorical values, collected from n samples. **(B-1) Identifying direct associations: causalDeepVASE incorporates** a statistical graphical model that selects interaction terms to identify associated variable pairs **(B-2)** An example of the identified direct associations (blue line). **(C-1) Identifying indirect associations: causalDeepVASE utilizes** deep neural network model to identify associated variable pairs interacting through mediating layers that are modeled with perceptron layers, which are indicated as orange nodes. After the first run sets V1 as response and identifies its association with other variables, causalDeepVASE will run this model with each of the other variables (V_2_, V_3_, V_m_) as response to identify all indirect associations. **(C-2)** An example of the identified direct associations (red line) **(D)** Incorporating the direct and indirect from both statistical learning model and deep learning model **(E)** Learning the causalities: causalDeepVASE utilizes degenerate gaussian (DG) to evaluate the causal direction for each association from **(C, D)**.

In the second step, after causalDeepVASE identifies the direct and indirect associations (**Figure 1B2, 1C2)**, it combines the associations (**Figure 1D**) and uses DG to learn causalities (**Figure 1E**) since this score criterion satisfies the two conditions to uniformly learn direct and indirect causalities, decomposability and flexibility. First, the DG score is proven to be decomposable under the assumption that the joint distribution over the independent variables factorizes according to the true causal structure such that a more parsimonious factorization does not exist. To be compatible with this assumption, which is the basis of several causal inference methods (4), causalDeepVASE removes variable associations that would violate this assumption, those that are conditionally dependent on any of the other variables (see Methods). Second, simulation studies showed that the DG score is flexible to identify the optimal causal structure that is generated outside of the assumed class. While the simulation studies showed this flexibility only using models of direct associations, the conditional Gaussian model and Lee and Hastie model (4), we confirmed that DG is flexible for indirect causalities in various simulated and biological data. Then, causalDeepVASE ensures to produce a directed acyclic graph (DAG) by manually removing the causal relations that create a cycle in reference to the effect size (see Methods). By using the DG score possessing decomposability and flexibility, causalDeepVASE learns direct and indirect causalities in a uniform and accurate fashion.

### causalDeepVASE identifies the true causal relations in simulated data

To assess the performance of causalDeepVASE, we will compare it to causalMGM, DG, and PC. Since DG learns causality based on the variable associations identified by an external method, we used the MGM graph using causalMGM’s implementation to learn causal relations with DG. Since the MGM implementation identifies the variable associations based on the interaction terms, we will refer to this model as the direct-DG model. Since causalDeepVASE runs DG on the direct and indirect associations, PC will be run on the direct and indirect associations to compare DG and PC. We will refer to this model as the direct-indirect-PC model (**Figure 2A**). Note that direct-DG and direct-indirect-PC models were developed in this paper for comparison purposes. In this section, we simulated data sets in the following way. With various sample sizes (n=200, 600, 1000), we simulated different numbers of variables (p = 50, 100, 200, 400, 600, 800, 1000, 1500, 2000, 2500, and 3000) where ten of the variables were selected to collectively determine the response value (see Methods). We simulated indirect causal relations to mimic complex biological mechanisms by implementing several unknown link functions between the ten true positives and the response (see Methods). We ran 20 repetitions in each simulation scenario. In the first step of identifying the variables associated with response, we compared four methods, causalDeepVASE, causalMGM, direct-DG, and direct-indirect-PC. For these methods, we reported the empirical power, which is the number of the true positives that are identified by each method. On average over the 20 repetitions, causalDeepVASE and the direct-indirect-PC outperform the other methods by identifying 80% of the true positives in most simulation scenarios, whereas causalMGM and the direct-DG consistently identified less than half of the true positives compared to causalDeepVASE and the direct-indirect-PC (**Figure 2B, Supplemental Table 1**).

**Figure 2.**
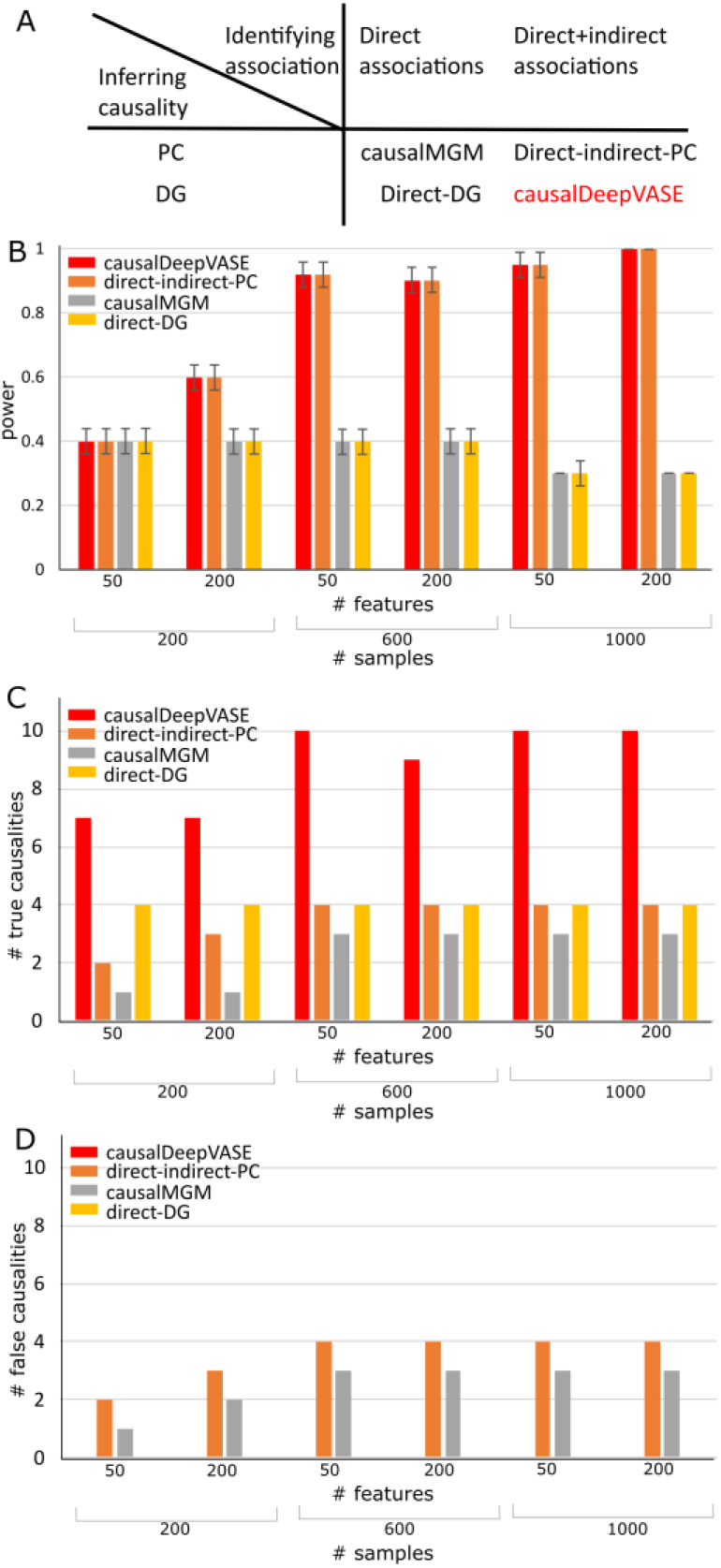
Performance assessment of four causal inference methods on the simulated data. **(A)** Combinations of association identifying modules (direct and indirect) and causality inference modules (DG and PC). **(B)** Average number of true associations identified by causalDeepVASE (red), direct-indirect-PC (orange), causalMGM (gray), or direct-DG (yellow) over 20 runs across various simulation scenarios, varying the number of features and the number of samples. The **(C)** Number of true causalities and **(D)** Number of false causalities identified by causalDeepVASE (blue), direct-indirect-PC (orange), causalMGM (gray), or direct-DG (yellow) at the 20^th^ run. CausalDeepVASE and direct-DG did not identify any false causalities.

In the second step where causalities are learned from the identified associations, we compared the number of true and false causalities (the 10 independent variables to response and response to the independent variables, respectively) of causalDeepVASE with those of the competing methods (**Supplemental Table 1**). In all scenarios, causalDeepVASE outperforms the other methods in identifying true causalities. Especially, for larger numbers of samples (n=600 and 1,000), causalDeepVASE identified most of the true causalities. On the other hand, causalMGM and direct-indirect-PC returned bidirectional causalities on the identified associations, which are counted as both true and false positives. Direct-DG learned the correct causalities on 4 associations (**Figure 2C**) it identified. Further, in distinguishing false-positive causalities, causalDeepVASE and direct-DG outperform other methods by returning 0 false-positive causalities, whereas causalMGM and indirect-direct-DG return some false positive causalities due to the bidirectional causalities (**Figure 2D**).

Altogether, causalDeepVASE outperforms other methods, identifying most of the true indirect associations and learning most causalities across various simulation scenarios without false positives, while competing methods could identify less than a half of true indirect causalities with some false positives.

### causalDeepVASE identifying indirect associations plays a critical role to understand complex biological systems

To understand why causalDeepVASE outperforms competing methods in identifying variable associations, we compared causalDeepVASE with causalMGM in identifying both direct and indirect associations on our pediatric sepsis data. We excluded direct-DG and direct-indirect-PC because they identify the same set of associations with causalMGM and causalDeepVASE, respectively. Our pediatric sepsis data (see Methods, n=283) consists of 56 demographic features (e.g. age, sex, health history) and cytokine variables (blood serum). Both causalDeepVASE and causalMGM identify 123 direct associations. An interesting finding among the direct associations is that serum level of soluble CD163 (sCD163) (19), a macrophage activator, associates with other biomarkers of macrophage activation, such as M-CSF (macrophage colony-stimulating factor) (20), MCP-1 (monocyte chemoattractant protein-1) (21), and other key drivers of macrophage response including hemoglobin (22) (**Figure 3A**). Also, heart rate and lymphocyte level (**Figure 3A**), whose normal ranges are defined based on age, are identified to have a direct association with age.

**Figure 3.**
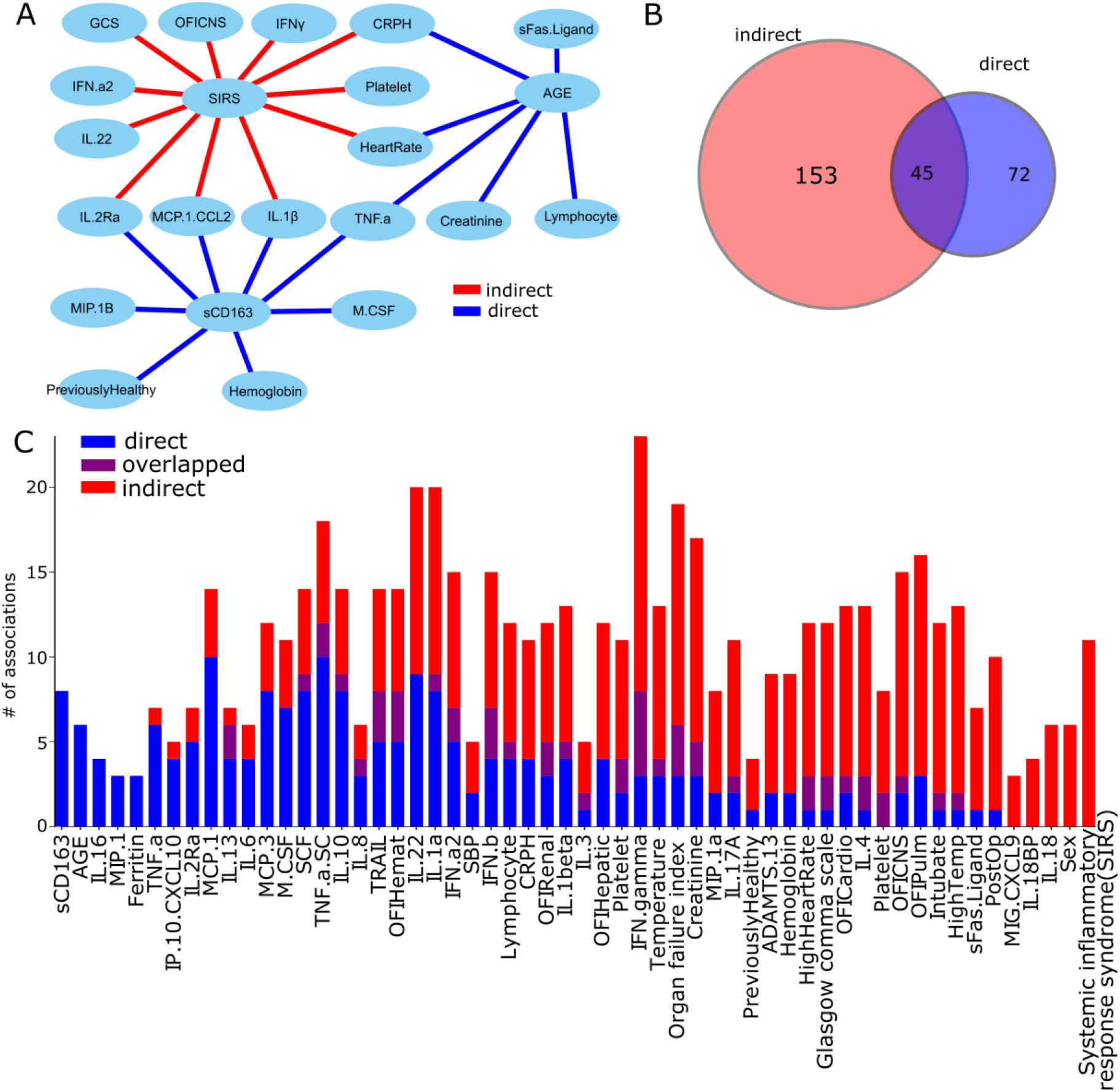
Direct and indirect associations in pediatric sepsis data. **(A)** Associations involving SIRS, AGE, and sCD163 identified as direct (blue) or indirect (red). **(B)** Ven diagram showing the intersection of feature associations, direct and indirect. While causalDeepVASE identified both direct and indirect associations, causalMGM identified only direct associations. **(C)** Number of associations involving each of the cytokines in the data.

In addition to this, causalDeepVASE identifies 201 indirect associations (**Supplemental Table 2**). After excluding the 48 associations that are identified also as direct associations, there are 153 unique indirect and 72 unique direct associations, identifying 2.1 times more indirect associations than direct associations (**Figure 3B**). Since the associations between cytokines are mostly mediated through multiple signal transduction pathways (23) (24), it is reasonable to identify more indirect associations than direct ones. Further inspection validated many of the indirect associations. Especially, we found that certain cytokines interact with other cytokines either only directly or indirectly (**Figure 3C**). For example, as the systemic inflammatory response syndrome (SIRS) status is clinically defined by four clinically recognizable criteria including elevated heart rate (25), SIRS is expected to be associated with heart rate. causalDeepVASE identified their association as indirect. Further, causalDeepVASE found indirect associations between SIRS and other cytokine features that are related to the immunologic state including IL-1β (26) and IFN-γ (27) (**Figure 3A**). Since SIRS represents immunological activation in tissues, this result together supports the value of identifying indirect associations.

By putting both direct and indirect associations together, causalDeepVASE allows us to look into the complex causal relationships, which would not be possible if considering only either direct or indirect associations. For example, the IFNγ levels have been suggested to interact with creatinine level (28) and causalDeepVASE can detect the interactions by incorporating direct and indirect associations (**Figure 3A**), whereas direct or indirect associations alone cannot connect creatinine to IFNγ. Altogether, in complex biological systems, biomedical variables may interact directly and/or indirectly. And causalDeepVASE enables the understanding of the complex biological systems by identifying both direct and indirect associations.

### causalDeepVASE identifies complex and non-linear associations validated between nutrients/gut bacteria and body-mass index (BMI)

In this section, we investigated how causalDeepVASE outperforms the other methods in learning true causalities by incorporating indirect causalities. We analyzed a cross-sectional data set consisting of 214 nutrient intakes as well as 87 bacteria genera in the gut and body-mass index (BMI) collected from 90 healthy volunteers. To evaluate the performance of causalDeepVASE, we investigated 16 indirect associations that were validated (13), 8 nutrient intakes and 8 bacteria genera. For these associations, causalDeepVASE and direct-indirect-PC outperform causalMGM and direct-DG again, where causalDeepVASE and direct-indirect-PC identified all 16 associations, while causalMGM and direct-DG identified 5 associations (31.3%) to BMI (**Figure 4A**). To further understand why the 11 associations were identified by causalDeepVASE and direct-indirect-PC, but not by causalMGM and direct-DG, we examined how 16 nutrient intake/bacteria genera levels change by the BMI values. The 5 associations identified by all the methods show a single linear association throughout the BMI region (**Figure 4B, S. Figure 1A-J**). On the other hand, the other 11 associations identified by causalDeepVASE and direct-indirect-PC show multiple sub-trends in various BMI ranges (**Figure 4C, S. Figure 1K-O**). For example, choline, phosphatidylcholine w/o suppl. intake (**Figure 4C**) shows an increasing trend from BMI 1~3, a decreasing trend from BMI 3~4, then another increasing trend from BMI 4~5. These multiple sub-trends may represent that mediating layers between the intake of choline, phosphatidylcholine w/o suppl. and the BMI status affect differently in different parts of the BMI region.

**Figure 4.**
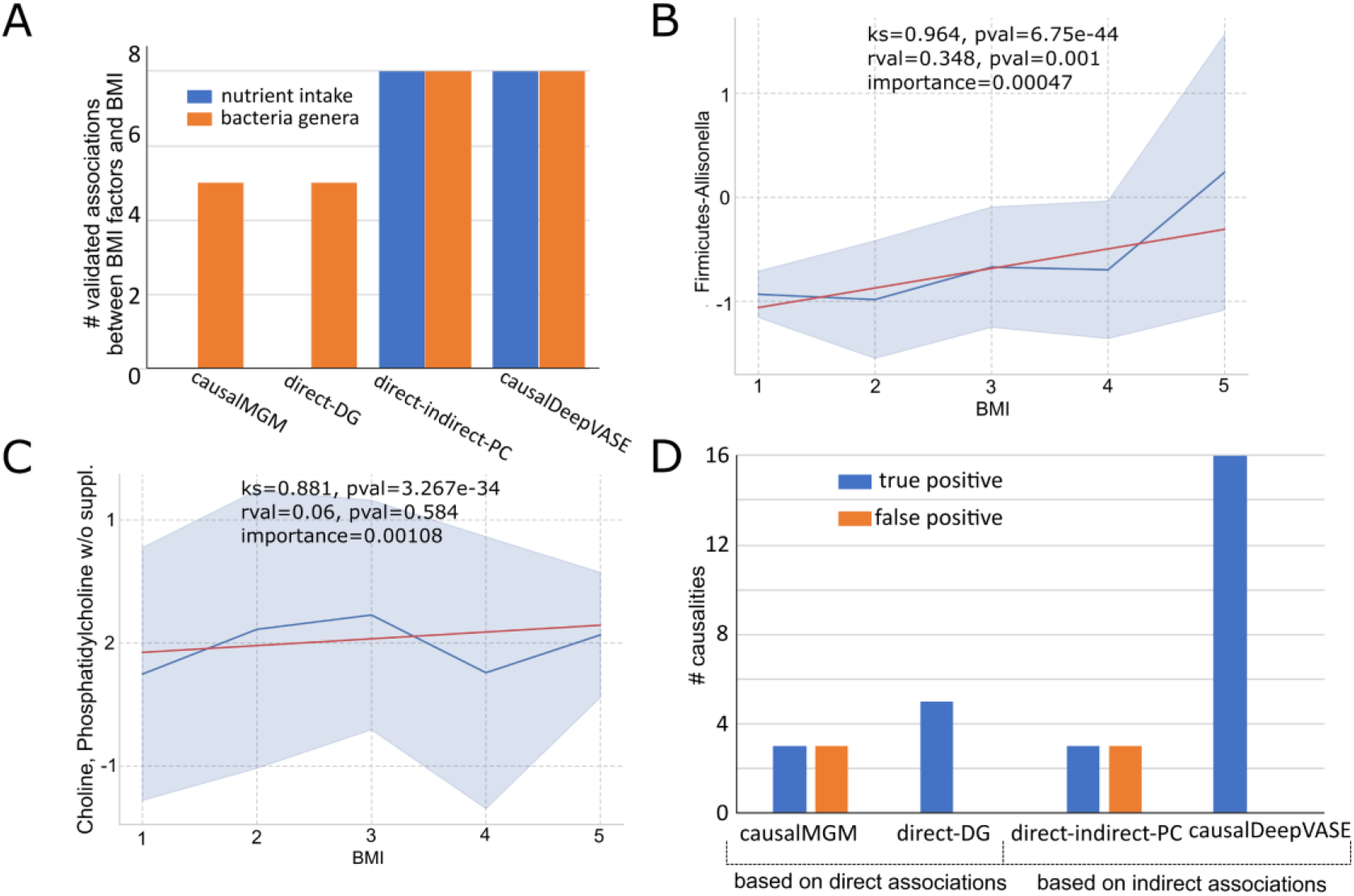
Performance assessment of four causal inference methods on BMI/bacteria/gut microbiome data. **(A)** Number of associations the methods (causalMGM, direct-DG, direct-indirect-PC, and causalDeepVASE) identified between the BMI status and 8 nutrient intake (blue) and 8 bacteria genera in the gut (red) that are validated associated with the BMI status. (**B**) Variable association relationship identified as direct association between BMI and Firmicutes-Allisonella. (**C**) Variable association relationship identified as indirect association between BMI and Choline, Phosphatidylcholine w/o suppl. In the relationship figures, KS is the KS test statistic, p-value is estimated from the KS test, rval is from a linear regression model, pval is from the linear regression, and importance is measured in causalDeepVASE. The gray indicates confidence intervals, the blue line indicates median values, and the red line represents the linearly regressed line. **(D)** Number of true causalities and false causalities identified by causalDeepVASE (blue), direct-indirect-PC (orange), causalMGM (gray), or direct-DG (yellow) at the 20th run. CausalDeepVASE and direct-DG did not identify any false causalities.

For the causalities from the associations, we deemed the causal direction from either nutrient intake or bacteria genera to BMI true. While causalDeepVASE identified the true causal direction in all the 16 identified associations, causalMGM and direct-indirect-PC identified 3 bidirectional causalities and direct-DG identified the true causal direction in the 5 identified associations (**Figure 4D**). Altogether, causalDeepVASE outperforms existing methods both in identifying associations (**Figure 4A**) and learning causalities (**Figure 4D)**, since it identifies and incorporates linear (**Figure 4B, S. Figure A-J**) and non-linear associations (**Figure 4C, S. Figure K-O**).

### causalDeepVASE identifies the causal relations across molecular and clinical variables

To understand how causalDeepVASE facilitates the understanding of complex biochemical mechanisms, we applied causalDeepVASE and the competing methods to learn causalities among gene expression and clinical variables of the TCGA breast cancer (29), including PAM50. PAM50 is an important clinical feature to categorize breast tumors with, which is defined by the expression information of 50 genes. To learn causalities from the genes to the PAM50 status, we considered 10 of the genes that are included in the 500 genes that are highly varying across 601 tumor samples in the data (high variance gene set). Therefore, the 10 genes are expected to have causal effects on the PAM50 status (PAM50-defining genes). We also considered 5 clinical variables, which are known to be associated with the PAM50 status: estrogen receptor (ER), human epidermal growth factor receptor (HER), and progesterone receptor (PER), lymph node status, and tumor staging code (see Methods).

In the first step of identifying associations, causalDeepVASE and direct-indirect-PC identified 9 associations out of 10 associations between PAM50-defining genes and PAM50, while causalMGM and direct-DG identified only 5 of them, increasing the power by 40% due to the identification of indirect associations (**Figure 5A, S. Figure 2A**). To examine the 9 associations, we estimated the effect size of the associations (see Methods). The effect size is significantly higher for the 9 PAM50-defining genes than the 491 other genes in the high variance set (P-val=0.027, **Figure 5B**), showing that this estimation is valid in identifying the known and thus strong associations. In terms of the clinical variables associated with PAM50, causalDeepVASE and direct-indirect-PC identified all 5 associations while causalMGM and direct-DG identified only one association (**Figure 5A, S. Figure 2B**), suggesting that the 4 associations identified only by causalDeepVASE and direct-indirect-PC are indirect associations. The high true positive rate of causalDeepVASE and direct-indirect-PC is partially related to its ability to identify both direct and indirect associations, unlike the other methods.

**Figure 5.**
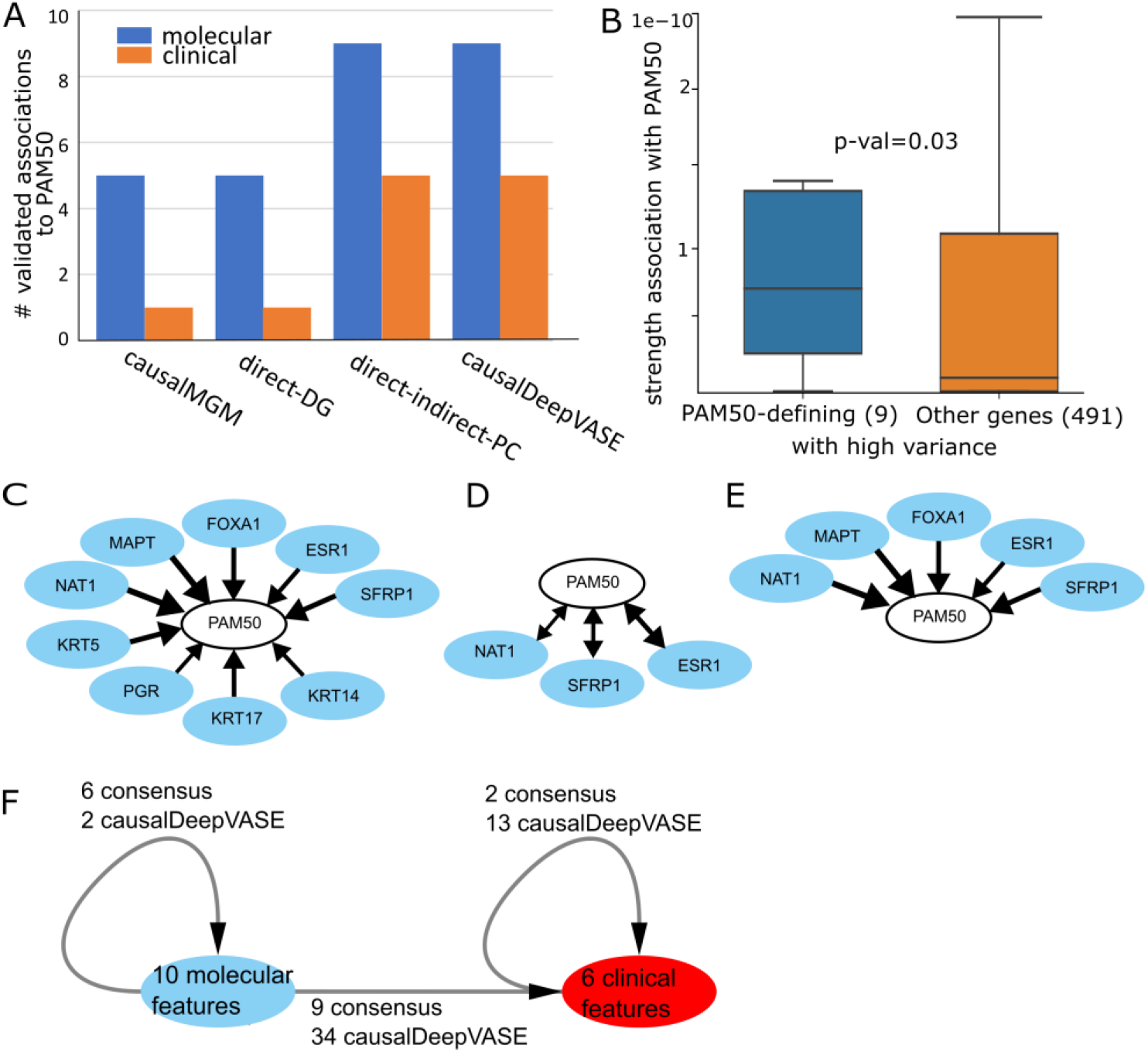
causalDeepVASE on TCGA breast cancer data. **(A)** Number of validated associations from molecular (blue) and clinical (orange) variables to PAM50. **(B)** Distribution of the association strength causalDeepVASE estimated for 9 PAM50-defining genes and 491 other genes in the high variance set. **(C, D, E)** Number of causalities identified by causalDeepVASE, causalMGM, DG in the default parameter settings, respectively. **(F)** Causalities over 10 molecular variables, 5 clinical variables, and the PAM50 status inferred by both the methods (causalMGM and causalDeepVASE) or by causalDeepVASE only.

In the second step of learning causalities from the identified associations in the first step, we compared the performance of the methods in various parameter settings (**Supplemental Table 3**). Between the PAM50-defining genes and the PAM50 status, we consider the inferred causalities from the genes to the PAM50 status as true positives since the PAM50-defining genes define the PAM50 status. causalDeepVASE outperforms both causalMGM and direct-DG, identifying true causalities from all 9 identified associations (**Figure 5C**). On the other hand, causalMGM and direct-indirect-PC identified bidirectional causalities for 3 associations and direct-DG identified the right causalities on all the 5 identified associations (**Figure 5D, E)**. We do not assess true and false positives in the clinical variables, since the causal direction is not clear in the clinical variables.

To examine how causalDeepVASE’s high sensitivity helps understand the complex breast tumor biology, we ran causalDeepVASE and causalMGM across the 10 molecular variables and the 6 clinical features, including PAM50 (**Figure 5F, S. Figure 2C**). Within the molecular variables, 2 of 8 (25%) causalities are identified uniquely by causalDeepVASE. However, within the clinical features and across molecular variables and clinical features, 13 out of 15 (86.7%) and 34 out of 43 (79.1%) causalities are identified uniquely by causalDeepVASE respectively, identifying 3-fold more causalities uniquely by causalDeepVASE. This high number of causalities within the clinical features and across molecular variables and clinical features is supported by recent studies demonstrating that the clinical features interact extensively both within and with the molecular variables in complex breast tumor biology (e.g., (30)(31)(32)(33)). Further, while the causalDeepVASE-unique causalities are likely indirect, their indirect relationships are supported by the fact that the clinical features are hormone receptor status and the PAM50 status, which are generated and regulated through multiple endocrine factors. Altogether, causalDeepVASE identifies indirect causalities across biological variables and clinical features in complex biological systems due to its sensitivity in identifying indirect associations and learning indirect causalities.

## DISCUSSION

In this paper, we developed one of the first computational methods, causalDeepVASE to learn both direct and indirect causalities in complex biological systems. It combines both statistical- and deep-learning models: the statistical-learning model identifies direct associations, and the deep-learning model identifies indirect associations. Applying causalDeepVASE to the simulated and biological data (pediatric sepsis, TCGA breast cancer, BMI with nutrients and gut bacteria) shows that causalDeepVASE outperforms existing methods (1) in identifying true associations because it incorporates indirect associations in the first step (**Fig. 1B, C**) and (2) in learning indirect causalities because of the decomposability and flexibility of the causal learning module, DG, in the second step (**Fig. 1D, E**). Due to the increased power, causalDeepVASE reveals a novel causal structure across molecular and clinical variables in complex biological systems. For example, causalDeepVASE uniquely identifies multiple causal relations across genes and clinical features (4.8 fold more than causalMGM) and within clinical features (7.5 fold more than causalMGM) in the TCGA breast cancer data (**Fig. 5F, S. Fig. 2C**).

In addition to the increased power, causalDeepVASE’s unique feature is to estimate effect size uniformly for both direct and indirect associations. Based on this estimate, we showed that the validated casualties are stronger than non-validated ones in both the breast cancer (P-value=0.03, **Fig. 5B**) and the BMI data (P-value=0.02, **S. Fig. 1P**). Assuming that validated associations are more apparent and stronger, these results suggest that causalDeepVASE’s effect size estimation successfully distinguishes strong associations from less strong ones, either direct or indirect.

causalDeepVASE is the first method that identified both direct and indirect causal relations. Especially, to learn causalities from direct and indirect associations, causalDeepVASE utilizes DG. DG was reported to perform well for learning direct causal relations when learning causal models, e.g., Lee and Hastie model, are generated outside of its assumed class (4), e.g., conditional Gaussian. With our experiments on identifying indirect causalities, we showed that DG outperforms the other established causal learning algorithm, PC (**Fig. 4D, 5F**), and identifies almost all true causalities.

In our data sets, causalMGM learned bidirectional causalities on most associations (**Fig. 4D, 5D**). Previous studies pointed out that the causal inference algorithm may return bidirectional causalities between two variables if there is a latent variable that causes both the variables^8^. Thus, collecting more data and thus including the latent variables may refine some of the bidirectional causalities into unidirectional causal relations.

In conclusion, we developed causalDeepVASE, which learns causal structure in complex biological systems. causalDeepVASE is one of the first methods to use a deep neural network approach to learn both direct and indirect causalities. Using causalDeepVASE, we found that complex biological systems have indirect causal relationships between molecular and clinical variables. In identifying such causal relationships, causalDeepVASE outperforms existing methods, causalMGM and DG. With these advantages, future application of causalDeepVASE can facilitate the identification of driver genes and therapeutic agents in biomedical studies and clinical trials.

## AVAILABILITY

The open-source causalDeepVASE program (version 0.9.1) is freely available at https://github.com/ZhenjiangFan/causalDeepVASE with necessary example data for this analysis.

## SUPPLEMENTARY DATA

**Supplementary Table 1.** Performance of the four methods (causalMGM, direct-DG, direct-indirect-PC, and causalDeepVASE) on the simulated data over various simulation settings

**Supplementary Table 2.** 201 Indirect feature associations identified by causalDeepVASE on our pediatric sepsis data

**Supplementary Table 3.** Performance of the four methods (causalMGM, direct-DG, direct-indirect-PC, and causalDeepVASE) on the breast cancer data over various parameter settings

## ACKNOWLEDGEMENT

We thank William Stafford Noble, Ph.D., Professor in Department of Genome Sciences and Department of Computer Science and Engineering, University of Washington for valuable discussion and their simulated and biological data. We thank Gregory Cooper, MD, Ph.D., Professor in Department of Biomedical Informatics, University of Pittsburgh for valuable discussion on causal structure learning. This research was supported in part by the University of Pittsburgh Center for Research Computing through the resources provided.

## TABLE AND FIGURES LEGENDS

**Supplementary Figure 1. (A-J)** Variable association relationship identified as indirect association between BMI and the feature indicated in the y-axis **(K-O)** Variable association relationship identified as direct association between BMI and the feature indicated in the y-axis. In the relationship figures, KS is the KS test statistic, p-value is estimated from the KS test, rval is from a linear regression model, pval is from the linear regression, and importance is measured in causalDeepVASE. The gray area indicates confidence intervals, the blue line indicates median values, and the red line represents regressed line. **(P)** Distribution of causalDeepVASE’s strength measure for the 16 validated factors and 285 other factors to BMI. **(Q)** K-S test statistic for 16 validated factors and 285 other factors to BMI **(R)** Distribution of the P-values from the K-S test estimated for 16 validated factors and 285 other factors to BMI.

**Supplementary Figure 2. (A)** PAM50-defining genes associated with the PAM50 status of patients identified by both causalDeepVASE and causalMGM (purple) or uniquely by causalDeepVASE (red). Both causalDeepVASE and causalMGM could not identify a PAM50-defining gene (MIA) in a dotted line. **(B)** clinical features (e.g. hormone status) of the breast cancer samples associated with PAM50. **(C)** Causalities inferred by causalDeepVASE over 10 molecular variables, 5 clinical variables, and the PAM50 status indicating how many causalities are inferred by both the methods (causalMGM and causalDeepVASE) or by causalDeepVASE only. ‘person_neoplasm_cancer_status’ refers to the state or condition of an individual’s neoplasm. ‘PR_status’, ‘ER_status’, and ‘HER2_status’ refer to the status of progesterone receptor, estrogen receptor, and human epidermal growth factor 2 receptor in the tumor sample.

## REFERENCES

1. Kim, S., Park, H.J., Cui, X. and Zhi, D. (2020) Collective effects of long-range DNA methylations predict gene expressions and estimate phenotypes in cancer. Sci. Rep., 10, 3920.

2. Kim, S., Bai, Y., Fan, Z., Diergaarde, B., Tseng, G.C. and Park, H.J. (2021) The microRNA target site landscape is a novel molecular feature associating alternative polyadenylation with immune evasion activity in breast cancer. Brief. Bioinform., 22.

3. Fan, Z., Kim, S., Bai, Y., Diergaarde, B. and Park, H.J. (2020) 3’-UTR Shortening Contributes to Subtype-Specific Cancer Growth by Breaking Stable ceRNA Crosstalk of Housekeeping Genes. Front. Bioeng. Biotechnol., 8, 334.

4. Andrews, B., Ramsey, J. and Cooper, G.F. (2019) Learning High-dimensional Directed Acyclic Graphs with Mixed Data-types. In Le, T.D., Li, J., Zhang, K., Cui, E.K.P., Hyvärinen, A. (eds), Proceedings of Machine Learning Research, Proceedings of Machine Learning Research. PMLR, Anchorage, Alaska, USA, Vol. 104, pp. 4–21.

5. Sedgewick, A.J., Shi, I., Donovan, R.M. and Benos, P. V (2016) Learning mixed graphical models with separate sparsity parameters and stability-based model selection. BMC Bioinformatics, 17, S175.

6. Lee, J. and Statistics,T.H.B.T.-P. of the S.I.C. on A.I. and Structure Learning of Mixed Graphical Models. 31, 388–396.

7. Spirtes, P. and Glymour, C. (1991) An Algorithm for Fast Recovery of Sparse Causal Graphs. Soc. Sci. Comput. Rev., 9, 62–72.

8. Raghu, V.K., Ramsey, J.D., Morris, A., Manatakis, D. V, Sprites, P., Chrysanthis, P.K., Glymour, C. and Benos, P. V (2018) Comparison of strategies for scalable causal discovery of latent variable models from mixed data. Int. J. data Sci. Anal., 6, 33–45.

9. Hasin, Y., Seldin, M. and Lusis, A. (2017) Multi-omics approaches to disease. Genome Biol., 18, 83.

10. Kim, S., Forno, E., Yan, Q., Jiang, Y., Zhang, R., Boutaoui, N., Acosta-Pérez, E., Canino, G., Chen, W. and Celedón, J.C. (2020) SNPs identified by GWAS affect asthma risk through DNA methylation and expression of cis-genes in airway epithelium. Eur. Respir. J., 55.

11. Sedgewick, A.J., Buschur, K., Shi, I., Ramsey, J.D., Raghu, V.K., Manatakis, D. V, Zhang, Y., Bon, J., Chandra, D., Karoleski, C., et al. (2019) Mixed graphical models for integrative causal analysis with application to chronic lung disease diagnosis and prognosis. Bioinformatics, 35, 1204–1212.

12. Candès, E., Fan, Y., Janson, L. and Lv, J. (2018) Panning for gold: ‘model-X’ knockoffs for high dimensional controlled variable selection. J. R. Stat. Soc. Ser. B (Statistical Methodol., 80, 551–577.

13. Lu, Y., Lv, J., Fan, Y. and Noble, W. (2018) DeepPINK: reproducible feature selection in deep neural networks.

14. Yu, Y., Chen, J., Gao, T. and Yu, M. (2019) {DAG}-{GNN}: {DAG} Structure Learning with Graph Neural Networks. In Chaudhuri, K., Salakhutdinov, R. (eds), Proceedings of the 36th International Conference on Machine Learning, Proceedings of Machine Learning Research. PMLR, Vol. 97, pp. 7154—7163.

15. Young, J., Andrews, B., Cooper, G. and Lu, X. (2020) Learning Latent Causal Structures with a Redundant Input Neural Network.

16. Carcillo, J.A., Berg, R.A., Wessel, D., Pollack, M., Meert, K., Hall, M., Newth, C., Lin, J.C., Doctor, A., Shanley, T., et al. (2019) A Multicenter Network Assessment of Three Inflammation Phenotypes in Pediatric Sepsis-Induced Multiple Organ Failure. Pediatr. Crit. care Med. a J. Soc. Crit. Care Med. World Fed. Pediatr. Intensive Crit. Care Soc., 20, 1137—1146.

17. Candes, E., Fan, Y., Janson, L. and Lv, J. (2016) Panning for Gold: Model-free Knockoffs for High-dimensional Controlled Variable Selection. J. R. Stat. Soc. Ser. B (Statistical Methodol., 80.

18. Chen, J. and Li, H. (2013) Variable selection for sparse Dirichlet-multinomial regression with an application to microbiome data analysis. Ann. Appl. Stat., 7, 418–442.

19. Crayne, C.B., Albeituni, S., Nichols, K.E. and Cron, R.Q. (2019) The Immunology of Macrophage Activation Syndrome. Front. Immunol., 10, 119.

20. Ushach, I. and Zlotnik, A. (2016) Biological role of granulocyte macrophage colony-stimulating factor (GM-CSF) and macrophage colony-stimulating factor (M-CSF) on cells of the myeloid lineage. J. Leukoc. Biol., 100, 481–489.

21. Deshmane, S.L., Kremlev, S., Amini, S. and Sawaya, B.E. (2009) Monocyte Chemoattractant Protein-1 (MCP-1): An Overview. J. Interf. Cytokine Res., 29, 313–326.

22. Finn, A. V, Nakano, M., Polavarapu, R., Karmali, V., Saeed, O., Zhao, X., Yazdani, S., Otsuka, F., Davis, T., Habib, A., et al. (2012) Hemoglobin directs macrophage differentiation and prevents foam cell formation in human atherosclerotic plaques. J. Am. Coll. Cardiol., 59, 166–177.

23. Stanley, A.C. and Lacy, P. (2010) Pathways for Cytokine Secretion. Physiology, 25, 218–229.

24. Leonard, W.J. and Lin, J.-X. (2000) Cytokine receptor signaling pathways. J. Allergy Clin. Immunol., 105, 877–888.

25. Merx, M.W. and Weber, C. (2007) Sepsis and the Heart. Circulation, 116, 793–802.

26. Nakanishi, K. (2018) Unique Action of Interleukin-18 on T Cells and Other Immune Cells. Front. Immunol., 9, 763.

27. Schoenborn, J.R. and Wilson, C.B. (2007) Regulation of interferon-gamma during innate and adaptive immune responses. Adv. Immunol., 96, 41–101.

28. Otto, G.P., Busch, M., Sossdorf, M. and Claus, R.A. (2013) Impact of sepsis-associated cytokine storm on plasma NGAL during acute kidney injury in a model of polymicrobial sepsis. Crit. Care, 17, 419.

29. Koboldt, D.C., Fulton, R.S., McLellan, M.D., Schmidt, H., Kalicki-Veizer, J., McMichael, J.F., Fulton, L.L., Dooling, D.J., Ding, L., Mardis, E.R., et al. (2012) Comprehensive molecular portraits of human breast tumours. Nature, 490, 61–70.

30. Angus, S.P., Stuhlmiller, T.J., Mehta, G., Bevill, S.M., Goulet, D.R., Olivares-Quintero, J.F., East, M.P., Tanioka, M., Zawistowski, J.S., Singh, D., et al. (2021) FOXA1 and adaptive response determinants to HER2 targeted therapy in TBCRC 036. npj Breast Cancer, 7, 51.

31. Onitilo, A.A., Engel, J.M., Greenlee, R.T. and Mukesh, B.N. (2009) Breast cancer subtypes based on ER/PR and Her2 expression: comparison of clinicopathologic features and survival. Clin. Med. Res., 7, 4–13.

32. Joly-Tonetti, N., Ondet, T., Monshouwer, M. and Stamatas, G.N. (2021) EGFR inhibitors switch keratinocytes from a proliferative to a differentiative phenotype affecting epidermal development and barrier function. BMC Cancer, 21, 5.

33. Ikeda, H., Taira, N., Hara, F., Fujita, T., Yamamoto, H., Soh, J., Toyooka, S., Nogami, T., Shien, T., Doihara, H., et al. (2010) The estrogen receptor influences microtubule-associated protein tau (MAPT) expression and the selective estrogen receptor inhibitor fulvestrant downregulates MAPT and increases the sensitivity to taxane in breast cancer cells. Breast Cancer Res., 12, R43–R43.

